# Interaction between decision-making and interoceptive representations of bodily arousal in frontal cortex

**DOI:** 10.1101/2021.02.04.429750

**Authors:** Atsushi Fujimoto, Elisabeth A. Murray, Peter H. Rudebeck

## Abstract

Decision-making and representations of arousal are intimately linked. Behavioral investigations have classically shown that either too little or too much bodily arousal is detrimental to decision-making, indicating that there is an inverted ‘U’ relationship between bodily arousal and performance. How these processes interact at the level of single neurons as well as the neural circuits involved are unclear. Here we recorded neural activity from orbitofrontal cortex (OFC) and dorsal anterior cingulate cortex (dACC) of macaque monkeys while they made reward-guided decisions. Heart rate (HR) was also recorded and used as a proxy for bodily arousal. Recordings were made both before and after subjects received excitotoxic lesions of the bilateral amygdala. In intact monkeys, higher HR facilitated reaction times (RTs). Concurrently, a set of neurons in OFC and dACC selectively encoded trial-by-trial variations in HR independent of reward value. After amygdala lesions, HR increased and the relationship between HR and RTs was reversed. Concurrent with this change, there was an increase in the proportion of dACC neurons encoding HR. Applying a novel population-coding analysis, we show that after bilateral amygdala lesions the balance of encoding in dACC is skewed away from signaling either reward value or choice direction towards HR coding around the time that choices are made. Taken together, the present results provide insight into how bodily arousal and decision-making are signaled in frontal cortex.

**Significance statement:** How bodily arousal states influence decision-making has been a central question in psychology, but the neural mechanisms are unclear. We recorded heart rate, a measure of bodily arousal, while simultaneously monitoring neural activity in orbitofrontal cortex (OFC) and dorsal anterior cingulate cortex (dACC) of macaques making reward-guided decisions. In intact macaques higher HR was associated with shorter reaction times. Concurrently, the activity of a set of neurons in OFC and dACC selectively encoded HR. Following amygdala lesions, HR generally increased and now the relationship between HR and reaction times was reversed. At the neural level, the balance of encoding in dACC shifted towards signaling HR, suggesting a specific mechanism through which bodily arousal influences decision-making.

## INTRODUCTION

Our current bodily state, whether it be a thirst or a racing heart, affects ongoing cognitive processes. Bodily arousal is fundamental to representations of our bodily state and can have a marked influence on decision-making (1–3). At moderate levels, bodily arousal can increase the chance of survival by invigorating responding, whereas at higher levels it promotes defensive behaviors such as freezing when threat of predation is imminent (4, 5). Consequently, altered generation and assessment of bodily arousal is thought to contribute to a host of psychiatric disorders such as anxiety disorders and addiction (6–8).

The influence of bodily arousal on behavior can be accounted for by viscerosensory feedback from the body reaching the brain. Central representations of current bodily state including arousal are known as *interoception* and these representations are thought to be essential for maintaining homeostasis (9). Several brain areas including the dorsal anterior cingulate cortex (dACC), orbitofrontal cortex (OFC), anterior insular cortex, hypothalamus, and amygdala are implicated in interoception, signaling bodily arousal as well as other aspects of physiological state such as hydration and temperature (10–13). The network of areas spanning frontal and limbic structures highlighted above as central to interoception overlaps extensively with the parts of the brain that are essential for reward-guided decision-making and these shared neural substrates are likely where these two processes interact (14, 15). Notably, lesions or dysfunction within frontal cortex and limbic areas in either humans or monkeys is associated with altered bodily arousal, interoception, and decision-making (16–18). Thus, optimal levels of bodily arousal are essential for appropriate responding to appetitive or aversive stimuli and likely require flexible adjustment of population-level neural representation in frontal and limbic structures (19).

Despite the appreciation that limbic and frontal structures are critical to both decision-making and interoception, how these processes interact in frontal cortex at the level of single neurons is poorly understood. This is because single neuron investigations of choice behavior have rarely considered or even attempted to measure the influence of bodily arousal on decision-making. Even less certain is how heightened states of bodily arousal affect interoceptive representations at the level of single neurons and subsequently influence choice behavior.

To address this, we analyzed a rare dataset: electrocardiogram (ECG) data was recorded simultaneously with single neuron recordings in OFC and dACC in macaque monkeys performing a reward-guided task both before and after excitotoxic lesions of the amygdala (20). In this previous study, we demonstrated that the decisions of monkeys were guided by the reward size associated with each option. In addition, we found that the reaction times (RTs) to choose rewarded options reflected the expected amount of reward. Correspondingly, the activity of a large proportion of single neurons in OFC and dACC preferentially encoded reward value. Here, our aim was to examine the potential interaction between factors that guide decision-making on a trial-by-trial basis (i.e. reward value and choice direction) and representations of bodily arousal in frontal cortex. For this purpose, we define heart rate as bodily arousal, and its neural representation as interoception. The heart rate during the fixation period of each trial (baseline HR) was used as a proxy of the current bodily arousal (21). Selective lesions of the amygdala caused a tonic increase in baseline HR which was seen both during reward-guided behavior (22) and at rest. Here we report that this increase in HR altered the influence of bodily arousal on decision-making, whereby heightened bodily arousal was associated with slower responding. At the same time, single neuron correlates of baseline HR increased in dACC, but not OFC, after amygdala lesions, altering the balance of coding away from decision-relevant processes and towards representations of bodily arousal. Taken together, this pattern of results suggests that bilateral amygdala lesions caused a state of hyper-arousal which impacts decision-making through adjustments in population coding in dACC.

## RESULTS

### Choice behavior

Three monkeys (monkey H, N, and V) performed a two-choice reward-guided task for fluid rewards (**Fig. 1A** and **B**). Behavior was assessed before and after bilateral excitotoxic amygdala lesions (**Fig. 1C**) and has been reported previously (20). In brief, all monkeys successfully (> 93%) chose the option that led to the greatest amount of reward on almost every trial. Importantly, performance was not impacted by bilateral excitotoxic amygdala lesions, which was by design as we wanted to ensure similar levels of performance before and after lesions (***SI Appendix***, **Fig. S1**).

**Figure 1.**
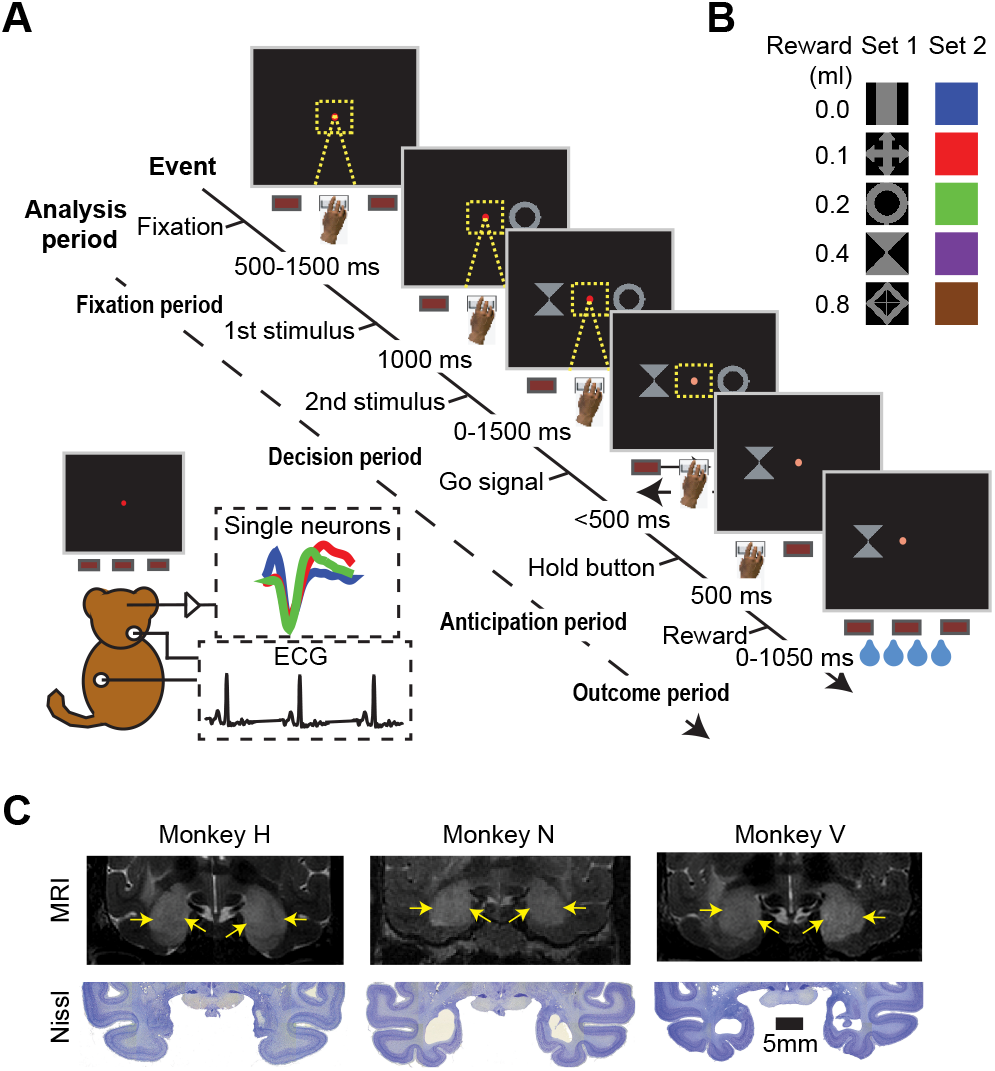
Reward-guided task and excitotoxic amygdala lesions. (*A*) The sequence of events in a single trial of the 2-option reward-guided task. Event timing and analysis periods are highlighted as well as dual neurophysiology and ECG recordings. A monkey could initiate a trial by placing its hand on a central button and fixing its gaze on a fixation spot in the center of the monitor. On each trial, monkeys were presented with two stimuli from a given stimulus set. After the fixation spot brightened (‘Go’ signal), the monkey could choose the left or right stimulus by moving its hand to the left or right button, respectively. After 500 ms, the reward amount assigned to the chosen stimulus was delivered. (*B*) The two stimulus sets used and corresponding reward amounts. (*C*) Bilateral excitotoxic lesions of the amygdala in monkeys H, N, and V as assessed by T2-weighted MRI scans taken within 5 days of infusions of excitotoxins (top row) and Nissl-stained sections at matching levels after histological processing (bottom row).

### Heart rate at rest and during a reward-guided task

Here we wanted to establish how the current state of bodily arousal influences decision-making. As an estimate of bodily arousal on a moment-to-moment basis in the task, we computed the heart rate during the fixation period (baseline HR) of each trial (21) (**Fig. 1A**). No stimuli were presented during this period; therefore, sensory and motor responses could not influence the baseline HR. This also ruled out the effect of the cardio-vascular reflex on HR associated with breath holding during liquid intake which might have obscured our results (23). Baseline HR was strongly correlated with HR in the other time periods analyzed (p < 0.01, rank-sum test, ***SI Appendix***, **Fig. S2A**), confirming that it is a robust predictor of bodily arousal across the whole trial.

As reported previously (22), the baseline HR significantly increased after bilateral amygdala lesions in all three monkeys (***SI Appendix***, **Fig. S2C**). This increase in baseline HR held both when monkeys were engaged in the behavioral task as well as at rest (p < 0.01, F_(1, 111)_ = 21.2, one-way ANOVA, ***SI Appendix***, **Fig. S2B**). This indicates that bilateral amygdala lesions caused a tonic increase in bodily arousal, regardless of whether the monkeys were engaged in a task or not. The increase in baseline HR is critical to interpreting the results that follow, as it indicates that effects on neural activity are associated with a general increase in bodily arousal.

### Interaction between reward-guided behavior and bodily arousal

Although the heightened bodily arousal following amygdala lesions did not disrupt choice accuracy, it might nevertheless have influenced choice behavior as indexed by reaction times (RTs). According to the Yerkes-Dodson law, moderate levels of bodily arousal should facilitate behavioral performance (24, 25). We thus reasoned that baseline HR should negatively correlate with RTs during the task in intact monkeys (**Fig. 2A**). To investigate this, we first compared the RTs in trials with higher baseline HR and compared them with those from trials with lower baseline HR using a median split. Before amygdala lesions, the difference in RTs tended to be lower than zero, indicating that the monkeys made faster response when the baseline HR was high (**Fig 2B**, left). To further quantify this effect, we performed a linear regression analysis with reward size, baseline HR, and their interaction term. There was a negative relationship between both reward size and baseline HR and RTs (rank-sum test, p < 0.01, df = 414), while the interaction between these variables was not significant (p = 0.72, df = 414) (**Fig. 2C**, left). Thus, both reward size and baseline HR independently facilitated RTs in intact monkeys, such that the monkeys responded faster when reward size was larger or baseline HR was higher.

**Figure 2.**
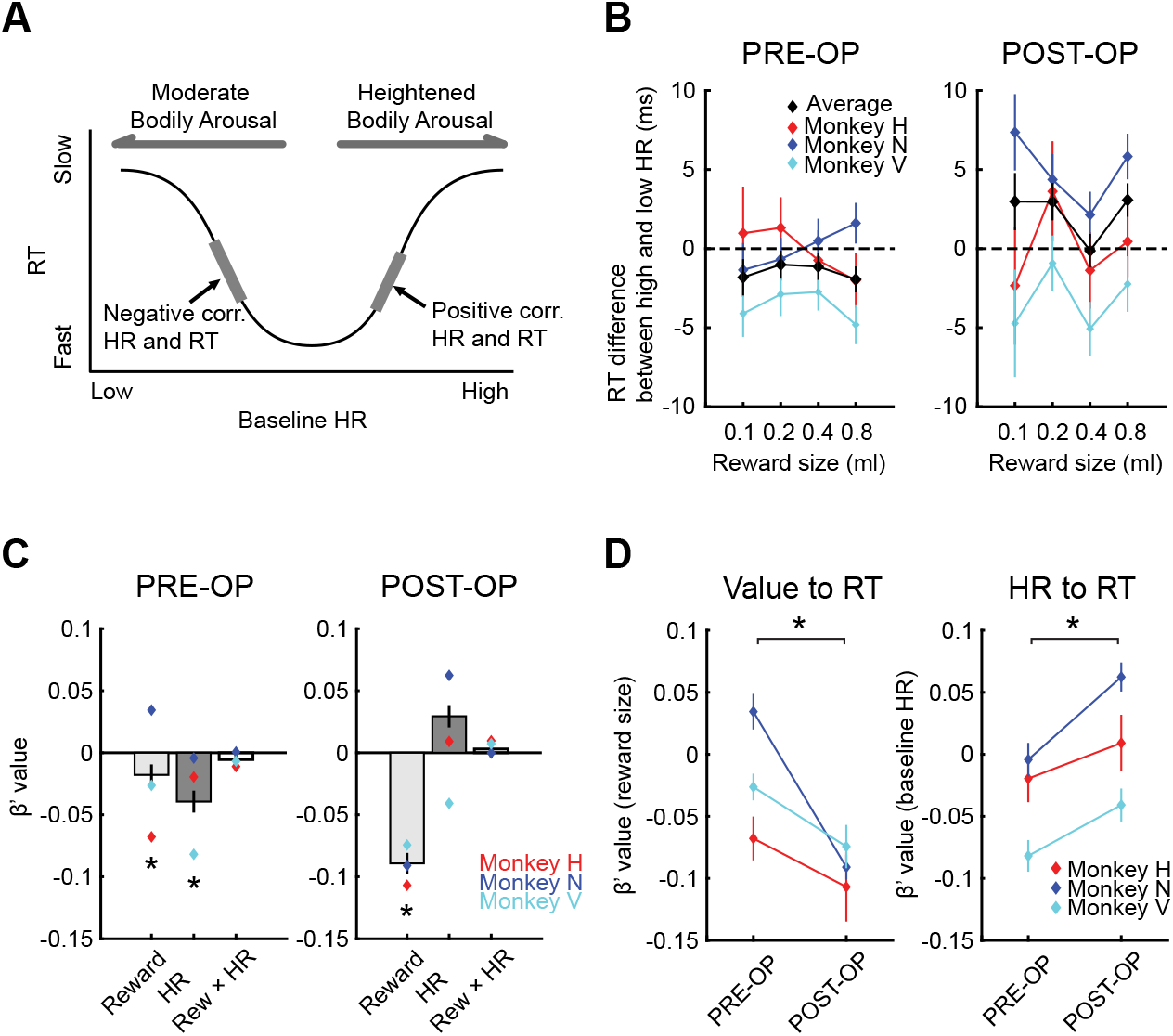
The influence of reward size, baseline HR, and their interaction on RTs. (*A*) Hypothetical relationship between baseline HR and RT. Bodily arousal facilitates behavioral performance when it is in the moderate range, whereas it inhibits performance when it is heightened. (*B*) Difference in RTs between trials with higher and lower baseline HR for each reward condition are plotted for pre- (left) and post-operative sessions (right). Positive value indicates larger RTs when baseline HR was higher. Red, blue, and cyan lines represent monkey H, N, and V, respectively, and black lines indicates the average of the three monkeys. (*C*) Average beta value from a multiple linear regression analysis of pre- (left) and post-operative (right) RTs. Diamond symbols represent the values for individual monkeys. Color convention as in *B*. Asterisks indicate statistical significance from zero (p < 0.05 with Bonferroni correction, rank-sum test). Error bars are standard error of the mean. (*D*) The impact of amygdala lesions on the influence of reward size (left) or baseline HR (right) on RTs is shown for each monkey. Asterisk indicates p < 0.05 by one-way ANOVA.

If higher HR is associated with faster RTs under normal conditions, we reasoned that tonically higher HR caused by amygdala lesions might have shifted subjects to the other side of the Yerkes-Dodson curve, such that baseline HR would positively correlate with RTs (**Fig. 2A**). Indeed, after amygdala lesions the difference in RTs between periods of high and low heart rate increased in all reward conditions, suggesting that increased HR had a broad impact on RTs (**Fig. 2B**, right). Two-way repeated-measures ANOVAs (amygdala lesions: pre-op or post-op × reward size:1, 2, 4, 8 drops) confirmed this as there was an effect of amygdala lesions (p = 0.037, F_(1,1659)_ = 4.4), but no effect of reward size or their interaction on RTs (p > 0.40).

We then conducted a multiple regression analysis to further investigate these effects. Notably, the relationship between baseline HR and RTs was reversed after lesions and the average beta value was now higher than zero (p = 0.039, df = 424, **Fig. 2C**, right) and the beta value for baseline HR increased in all monkeys after lesions (p < 0.01, F_(1, 415)_ = 11.8, **Fig. 2D**, right). In other words, now higher HR was associated with slower RTs. The effect of reward size on RTs was maintained (p < 0.01, df = 424, **Fig. 2C**, right) and, if anything, was greater after lesions (p < 0.01, F_(1, 415)_ = 30.8, **Fig. 2D**, left). Importantly, the direction of these effects was consistent across all the monkeys. Thus, bilateral amygdala lesions reversed the impact of arousal and concurrently enhanced the influence of reward value on RTs.

One possible explanation for this pattern of results is that baseline HR is directedly related to reward history. However, baseline HR was not associated with reward size in the preceding trial, either before or after amygdala lesions (***SI Appendix***, **Fig. S2C**, repeated-measures ANOVAs effect of lesion, p < 0.01, F_(1, 1660)_ = 346.8; effect of reward, p = 1.00, F_(3, 1660)_ = 0.015; lesion by reward interaction, p = 0.99, F_(3, 1660)_ = 0.025). In addition, baseline HR was not related to the average reward from the preceding trials (1-10 trials). The mean correlation coefficient (Pearson’s *r*) between baseline HR and average reward size did not significantly deviate from zero in sessions before (p = 0.28, n = 208, rank-sum test) or after amygdala lesions (p = 0.93, n = 213, ***SI Appendix***, **Fig. S2D**). Thus, the trial-by-trial fluctuations in the baseline HR were independent of rewards received in the prior trials.

We also tested whether the impact of amygdala lesions on RTs was influenced by choice direction. Two-way repeated-measures ANOVAs (amygdala lesions: pre-op or postop × choice direction: left or right) revealed no significant main effect of choice direction (p = 0.72, F_(1,830)_ = 0.11) or its interaction to amygdala lesions (p = 0.30, F_(1,830)_ = 1.1), suggesting that the impact of amygdala lesions on RTs was not influenced by the direction of choices.

### OFC and dACC neurons encode decision-making variables and current bodily arousal

In the present task, reward value was the critical factor that monkeys used to guide their choices, but on each trial the option associated with the greatest amount of reward could either be on the left or right side of the screen. Thus, once monkeys identified the best option they had to decide to choose left or right. Consequently, our analyses of how decision-making and bodily arousal interacted concentrated on how neuronal representations of reward value, choice direction, and bodily arousal interact when decisions are made. Prior studies have reported that neurons in OFC and dACC encode various aspects of reward, especially reward magnitude (26–28). How reward representations in frontal cortex interact with the current state of bodily arousal is unknown. Indeed, it is possible that encoding of reward magnitude or reward probability is confounded with bodily arousal, in that bodily arousal is the primary variable encoded and this has been misinterpreted as reflecting reward value. An alternative possibility is that potential rewards and arousal states are encoded separately by neurons in frontal cortex.

To address this question, we analyzed the activity of 271 OFC (31, 105, 135 neurons from monkeys H, N, and V, respectively) and 232 dACC neurons (102, 117, 13 neurons from monkeys H, N, and V, respectively) recorded before amygdala lesions (**Fig. 3A** and **B**). Here we included both well-isolated neurons and multi-unit activity in our analyses (see ***SI Appendix***, **Table S1** for a full breakdown). All results reported here hold irrespective of the inclusion of multi-unit activity.

**Figure 3.**
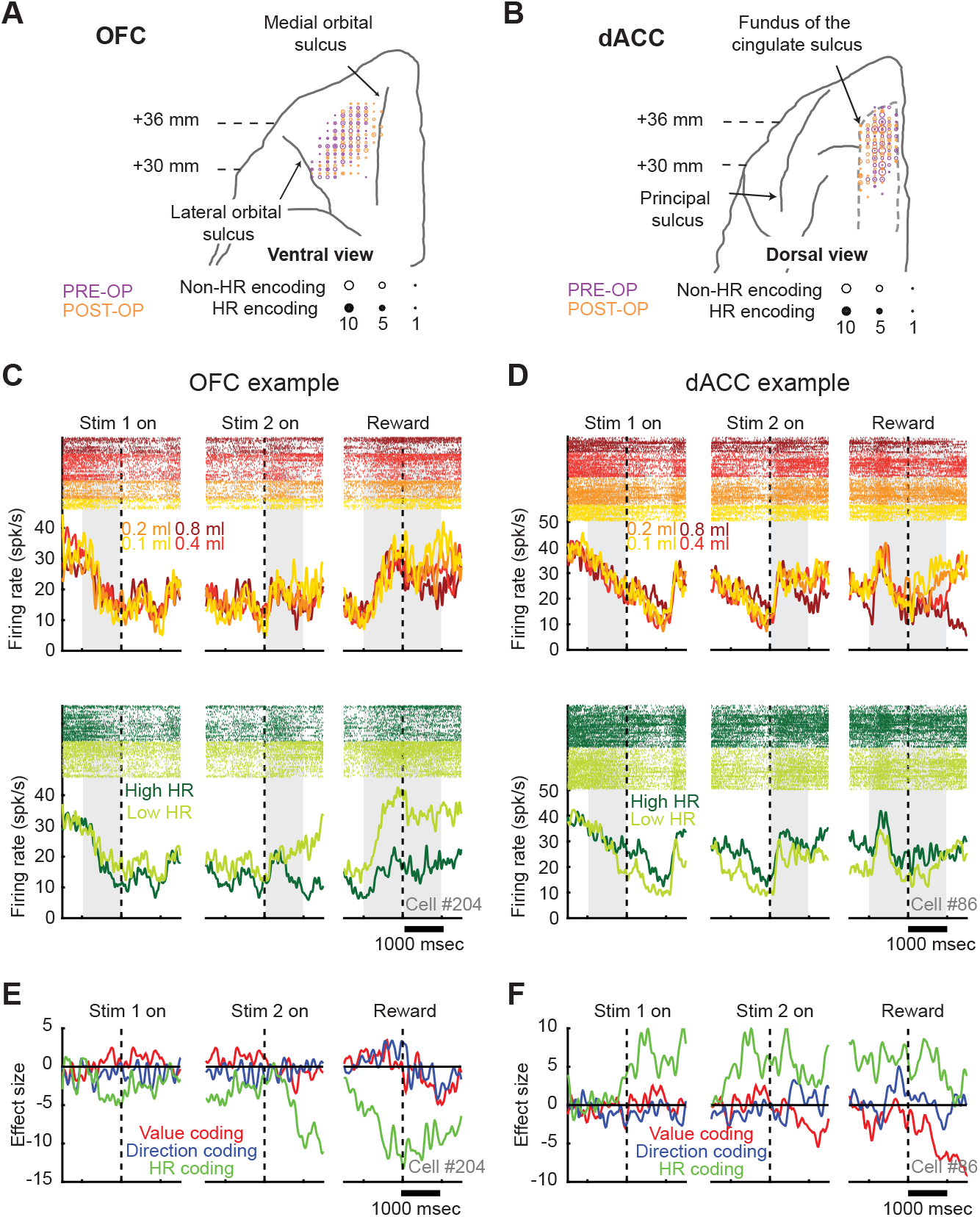
Encoding of baseline heart rate by neurons in OFC and dACC. (*A and B*) Schematics of recording locations in OFC (left) and dACC (right). Recording locations are plotted on the ventral view (OFC) or dorsal view (dACC) of a standard macaque brain. A-P distances (in mm) from the ear canal are shown. Larger symbols indicate increasing numbers of HR-coding neurons in the decision period. (*C and D*) Examples of HR coding neurons in OFC (*C*) and dACC (*D*). To visualize neuronal modulation by baseline HR, firing rate is plotted separately for those trials with low baseline HR (Low HR) and high baseline HR (High HR). Top and bottom panels show raster plots and peri-stimulus time histograms (PSTHs) aligned to the timing of onset of 1^st^ stimulus (left), onset of 2^nd^ stimulus (center), and reward delivery (right). Colors indicate the reward condition (top) and baseline HR (bottom). Gray areas indicate the 1,000-ms analysis time periods; from left to right, the analysis periods correspond to fixation, decision, anticipation and outcome. (*E and F*) Time course of beta coefficients from multiple linear regression including reward value (red), direction (blue), and baseline HR (green) of the neurons in *C* and *D*, respectively.

Figure 3C depicts an example OFC neuron that reflected baseline HR in its firing rate (HR coding). Here, to visualize modulation of neuronal activity by baseline HR, we divided trials into ‘high baseline HR’ and ‘low baseline HR’ using a median split. This neuron increased its activity when baseline HR was low, especially around the time when reward was delivered (**Fig. 3C**, bottom row). This neuron was defined as a ‘negative HR coding’ as its activity showed negative correlation to baseline HR during anticipation and outcome periods (p < 0.05, multiple linear regression). The same neuron was not strongly modulated by reward value during the same period (**Fig. 3C**, top row). Figure 3D depicts a neuron in dACC that had higher activity when baseline HR was high (**Fig. 3D**, bottom row). This neuron was defined as a ‘positive HR coding’ as its activity showed a positive correlation with baseline HR during the analysis epochs (p < 0.05). Notably, the same neuron exhibited negative reward-value coding during the outcome period (**Fig. 3D**, top row). Note that for both reward value and HR, the value of that variable is fixed within a trial (number of drops of reward, baseline HR) and the plots show modulation of firing rate by that variable across the trial. To assess the time course of encoding, a sliding window multiple linear-regression (100-ms bin shifted in 10-ms steps), in which firing rate of each neuron was modeled with reward size, choice direction, and baseline HR was conducted (Eq. 2, see Methods), for the same example neurons (**Fig. 3E** and **F**). This revealed that the baseline HR was encoded through the trial at a single-neuron level and was not always coupled to reward size or choice direction coding.

In the overall population of recorded neurons, the activity of a substantial number of neurons reflected the current level of bodily arousal as well as variables directly related to the decision-making process. During the decision period, reward value and choice direction were encoded by 35.4% and 10.7% of OFC and 28.0% and 22.0% of dACC neurons, respectively (**Fig. 4A**). In this period, OFC tended to encode reward value more strongly than dACC (96/271 and 65/232 in OFC and dACC, respectively; p = 0.076, χ^2^ = 3.2, chi-square test), while dACC encoded choice direction more strongly than OFC (29/271 and 51/232 in OFC and dACC, respectively; p < 0.01, χ^2^ = 11.9). During the same period, 15.5% of OFC (42/271) and the same proportion of dACC (36/232) neurons showed HR coding (p < 0.05, **Fig. 4A**). The proportion of HR coding was consistent across each monkey tested (p > 0.53, two-sample Kolmogorov-Smirnov test).

**Figure 4.**
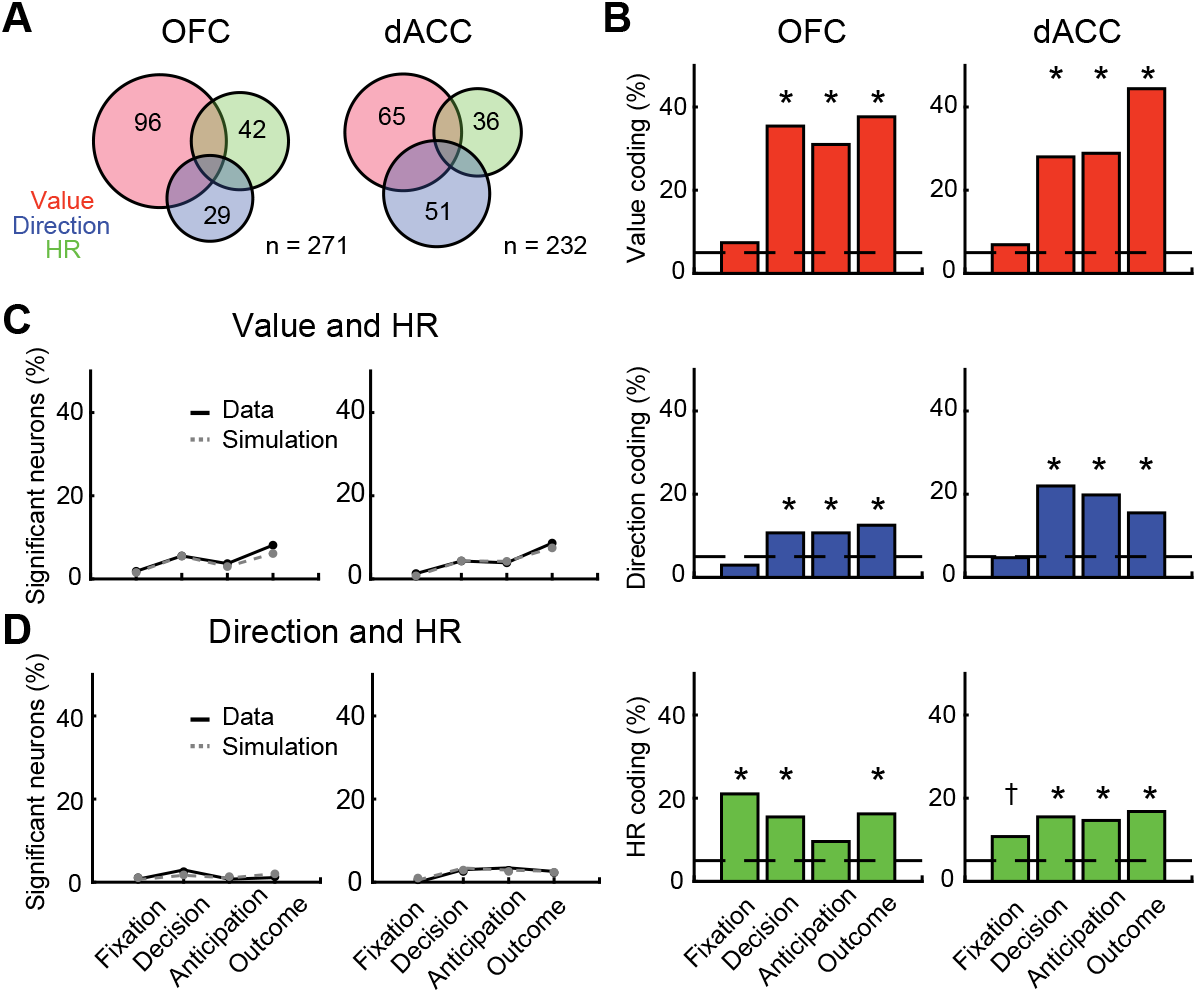
Proportion of neurons showing significant encoding before amygdala lesions. (*A*) The numbers of neurons showing significant encoding of reward value (size), choice direction, and HR in the decision period in OFC (left) and dACC (right), respectively. (*B*) Percentage of significant reward-value coding neurons (top), direction coding neurons (middle), and HR coding neurons (bottom) in each time period in OFC (left panels) and dACC (right panels). Dashed lines indicate chance level (5%). Asterisks indicate significant difference from chance level (* p < 0.05, † p < 0.10 with Bonferroni correction, chi-square test). (*C*) Percentage of neurons that showed both reward-value and HR coding at the same epoch (black plot) and simulated chance level (gray plot). (*D*) Percentage of neurons that showed both choice direction and HR coding. No significant difference between the data and simulation was observed in any analysis epoch by chi-square test (p > 0.38).

To address the dynamics of reward value, direction, and HR coding during a trial, we computed the proportion of significant neurons for each epoch (p < 0.05, **Fig. 4B**). As reported previously, more OFC and dACC neurons encoded reward value than choice direction (20). Although reward-value coding tended to be stronger in OFC during the decision period, no significant difference across areas was observed during anticipation and outcome periods (p > 0.10, chi-square test) (**Fig. 4B**, top row). Choice direction was preferentially encoded in dACC, and the difference across areas was pronounced during the decision and anticipation periods (p < 0.01, chi-square test) (**Fig. 4B**, middle row). These results highlight the different roles of OFC and dACC in the decision-making process, in which the former represents reward expectation and outcome and the latter represents choice direction as well as reward.

Encoding of HR was found in a substantial proportion of OFC and ACC neurons. In the majority of analysis windows, the proportion of neurons significantly exceeded chance in both areas (p < 0.05 with Bonferroni correction, chi-square test) chi-(**Fig. 4B**, bottom row).

While overall HR coding was indistinguishable between areas, HR coding during the fixation period was greater in OFC than in dACC (p = 0.0018, χ^2^ = 9.8, chi-chi-square test). A full description of the proportion of neurons encoding reward value, choice direction, and baseline HR is presented in ***SI Appendix***, **Table S2**.

To directly assess whether or not there was an interaction between the encoding of bodily arousal and reward, or, separately, between the encoding of bodily arousal and choice direction, at the level of single neurons, we first looked at the proportion of neurons that encoded both reward size and baseline HR during the same analysis window. The chance level was computed by multiplying the proportion of reward-value and HR coding neurons. The number of neurons encoding both factors did not exceed the chance level in either area (**Fig. 4C**, p > 0.41, chi-square test). Next, we tested the proportion of neurons that encoded both choice direction and baseline HR during the same analysis window. Again, the number of neurons encoding both factors did not exceed the chance level in either area (**Fig. 4D**, p > 0.38). These analyses indicate that the representation of current bodily arousal, choice direction, and reward value are encoded by largely independent populations of neurons in OFC and dACC.

### Bilateral amygdala lesions increase the proportion of HR coding neurons in dACC

Our analysis of HR demonstrated that bilateral lesions of amygdala increased the general level of bodily arousal and altered the relationship between bodily arousal and reaction time. Here we looked for how this change in bodily arousal might have altered encoding of HR and its interaction with reward value and choice direction. Following bilateral excitotoxic lesions of the amygdala we recorded the activity of 239 neurons in OFC (142, 97 neurons from monkeys N and V, respectively) and 190 in dACC (83, 107 neurons from monkeys H and N respectively). Recordings were conducted using identical procedures to those used before amygdala lesions and recording locations overlapped with those targeted before amygdala lesions (**Fig. 3A** and **B**).

Following amygdala lesions, the proportion of neurons encoding baseline HR was unchanged in OFC (average across all epochs, 15.6% to 15.2%), but was increased in dACC (average across all epochs, 14.4% to 21.3%) (**Fig. 5C**). Two-way repeated-measures ANOVAs with factors of lesion (pre-op, post-op) and area (OFC, dACC) revealed a significant interaction of lesion by area (p < 0.01, F_(1,5188)_ = 13.3), indicating that amygdala lesions altered the balance of encoding of HR across areas. This effect was especially pronounced in the fixation period where the proportion of dACC neurons encoding baseline HR increased by approximately 12% (10.8% to 22.6%, p = 0.0016, χ^2^ = 10.9) (**Fig. 5C**, center).

**Figure 5.**
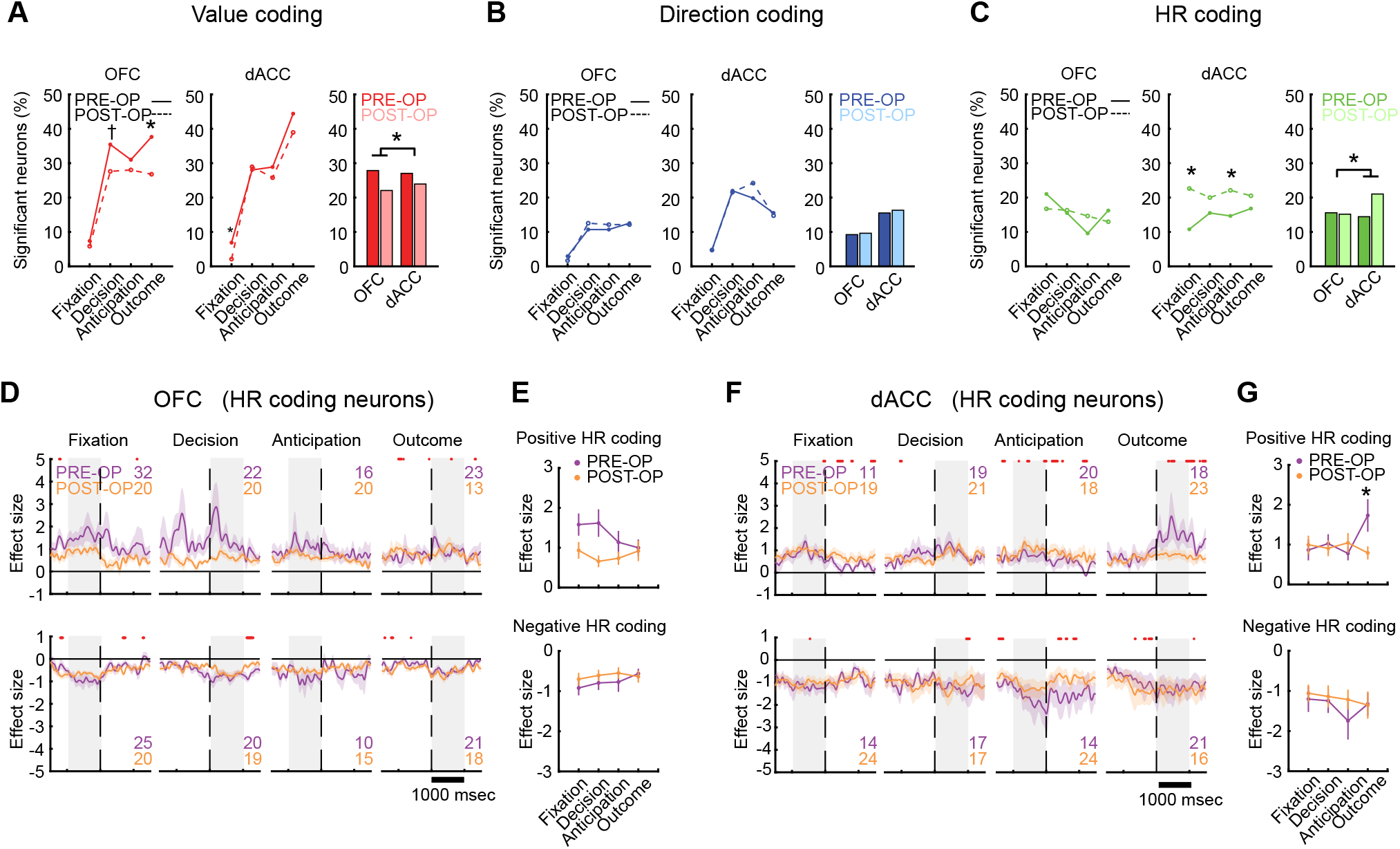
Effect of bilateral amygdala lesions on the proportion of neurons encoding reward amount, choice direction, and baseline HR. (*A*) *left and center*: Proportion of neurons that encoded the reward value in OFC (left) and dACC (center) in each of the four epochs analyzed. Solid lines represent pre-op and dashed lines represent post-op data. Asterisks indicate significant difference in the proportion of neurons for each coding type in each analysis period (p < 0.05, chi-square test). *right*: The average of proportion in the four epochs. Darker color represents pre-op and lighter color represents post-op data. Asterisk indicates significant interaction of lesion by area (p < 0.05, repeated-measures ANOVAs). (*B and C*) Proportion of direction coding neurons (*B*) and HR coding neurons (*C*) in OFC and dACC. Conventions as in *A*. (*D and F*) The time-course of effect size for HR coding in OFC (*D*) and dACC neurons (*F*) respectively. Positive HR coding (top row) and negative HR coding neurons (bottom row) are shown separately for each analysis epoch. The numerals in the panels indicate the number of HR coding neurons for each analysis epoch (purple: pre-op, orange: post-op). Red dots at the top of panels indicate significant difference in effect size between pre- and post-operative data (p < 0.05, rank-sum test) (*E and G*) Average effect size for each analysis period for OFC (*E*) and dACC (*G*). Asterisk indicates p < 0.05, rank-sum test.

We also performed the same analysis on reward-value coding and choice-direction coding. We found a significant interaction of lesion by area in reward-value coding (p = 0.034, F_(1,5188)_ = 4.5, **Fig. 5A**), but not in choice-direction coding (p = 0.90, F_(1,5188)_ = 0.016, **Fig. 5B**), which indicates that the amygdala lesions preferentially decreased reward-value coding in OFC as was reported in our prior study (20). These results held when we used the more conservative threshold of p < 0.0125 for the detection of significant neurons (***SI Appendix***, **Fig. S3**). Thus, following amygdala lesions which heightened bodily arousal in all three monkeys, there was an increase in the proportion of neurons encoding bodily arousal that was specific to the dACC.

We further investigated how representations of bodily arousal changed at the level of single neurons by looking at how amygdala lesions altered the strength of HR coding (i.e., the effect size) over time (**Fig. 5D-G**). Here we computed the time course of beta coefficients for baseline HR (Eq. 2, see Methods). Positive and negative HR-coding neurons, defined by their sign of the beta coefficient, were analyzed separately. In OFC, HR coding was lower during the fixation, decision, and anticipation periods following lesions (**Fig. 5D** and **E**). In dACC, HR coding significantly decreased during the outcome period but was not significantly altered in other periods (**Fig. 5F** and **G**). Thus, whereas amygdala lesions led to a selective increase in the proportion of dACC neurons encoding the current bodily arousal, this was not associated with an increase in the strength of encoding in single neuron activity.

### Contrasting effects of amygdala lesions on the population coding of reward, direction, and bodily arousal in OFC and dACC

Changes in the proportion of HR and reward-value coding neurons in OFC and dACC after amygdala lesions (**Fig. 5A-C**) could reflect a broader change in the balance of coding within these areas at the population level. To investigate this, we performed a population coding analysis on all of the neurons recorded both before and after lesions. We projected the beta coefficients from the regression analyses for reward value and HR coding for all recorded neurons, and fitted an ellipse representing the 95% confidence interval of the data (population ellipse).

If the coding in the neural population is generally decreased across all variables, then there should be a general reduction in the size of the ellipse (**Fig. 6A**). By contrast, if there is a rotation of the ellipse this would indicate a change in how the population of neurons is encoding the two variables (**Fig. 6B**). The analysis was performed for each of the decision, anticipation, and outcome periods. Before amygdala lesions, population ellipses were stretched along the horizontal axis in both areas, reflecting the prominence of reward-value relative to HR coding in both OFC and dACC (**Fig. 6C** and **E**, left columns).

**Figure 6.**
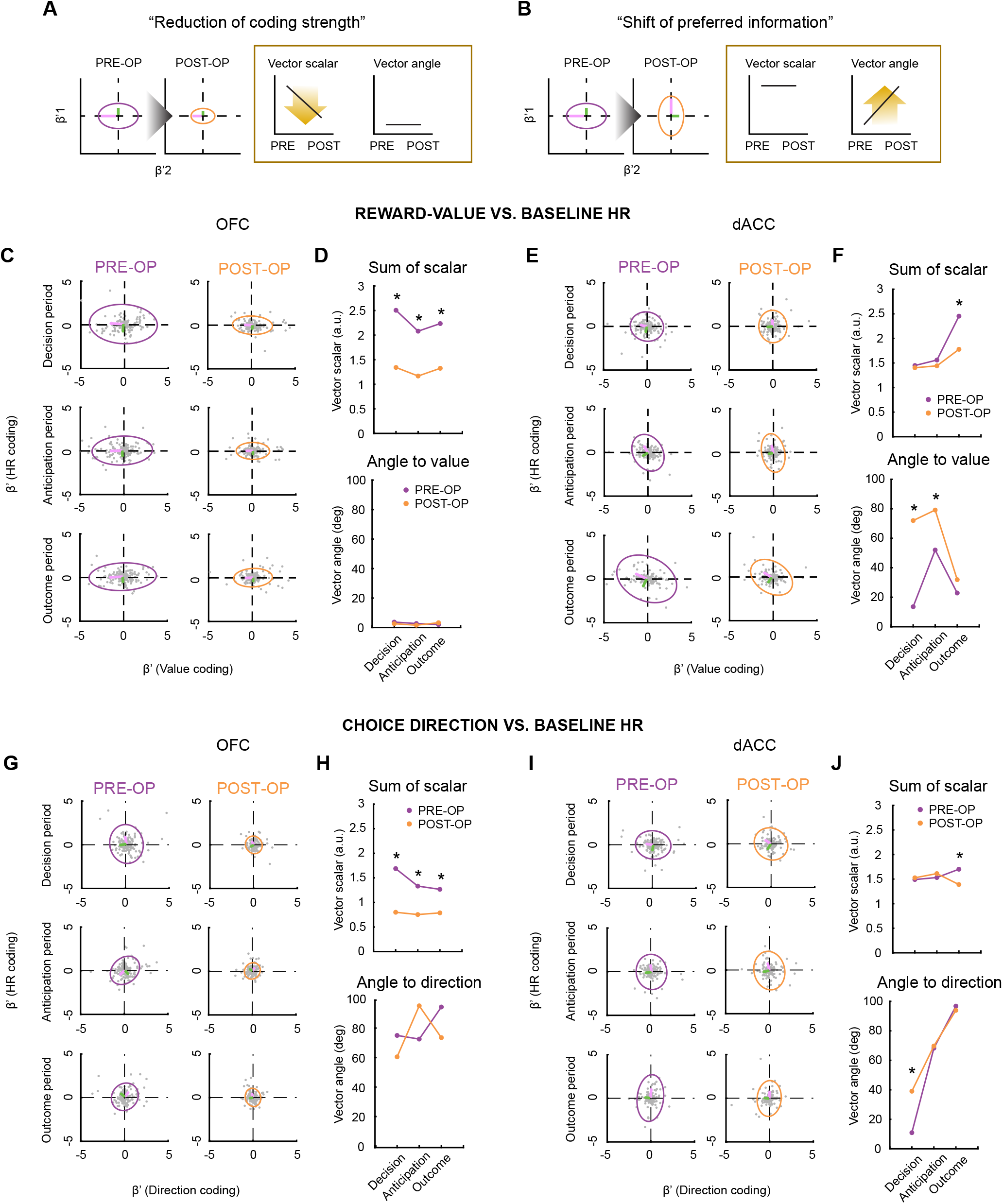
Effect of amygdala lesions on population representations of reward-value and baseline HR representations in OFC and dACC. (*A*) Schematic figure to illustrate how a general reduction of information coding would impact the scalar of the ellipse fitted to the encoding of the two variables. After lesions, a reduction in the size of the population ellipse (95% CI of the data) reflects a reduction of both reward-value and HR coding (left panels). The change can be projected to the reduction in the vector scalar but not the vector angle (right panels). (*B*) Schematic figure to illustrate how a shift in the preferred information would impact the vector angle of the population code. After lesions, a rotation of the population ellipse around the center reflects a reduction of reward-value coding and an increase of HR coding (left panels). This change can be projected to the change of vector angle but not the vector scalar (right panels). (*C*) Population representations of baseline HR and reward value in OFC neurons before (left column) and after (right column) amygdala lesions, for decision (top), anticipation (middle), and outcome (bottom) periods. Ellipse represents 95% confidence interval of the data (purple: pre-op, orange: post-op). Long (magenta) and short (green) eigenvectors are superimposed. (*D*) Vector scalar (top) and angle (bottom) change in OFC as function of amygdala lesion. (*E and F*) Population representation of baseline HR and reward value in dACC neurons. Conventions as in *C* and *D*. (*G-J*) Population representations of baseline HR and choice direction in OFC neurons (*G* and *H*) and dACC (*I* and *J*). Conventions as in *C-F*. Asterisks indicate AUC > 0.90.

The bilateral amygdala lesions differently affected the OFC and dACC populationcoding patterns. In the OFC, population ellipses tended to shrink after amygdala lesions, reflecting a decrease in encoding of both reward value and baseline HR (**Fig. 6C**, right column). In the dACC, on the other hand, the population ellipses maintained their shape, but tended to rotate around the center, reflecting an increase of HR coding and a decrease in reward-value coding (**Fig. 6E**, right column).

To quantify these changes, we computed the sum of scalar of the long and short eigenvectors, and the angle between the long eigenvector and the value-coding axis, for randomly subsampled population data (see **Fig. 6A** and **B**). We assumed that the vector scalar represents the strength of information coding, and the vector angle represents the relative weighting of the strength between variables. In OFC, bilateral amygdala lesions exclusively affected vector scalar (**Fig. 6D**, top). The sum of the scalar decreased during decision, anticipation and outcome periods (AUC > 0.90), reflecting the reduction in both reward-value and HR coding. No corresponding change in vector angle was apparent (**Fig. 6D**, bottom). By contrast, in dACC, amygdala lesions affected the vector angle and this change was most prominent during the decision and anticipation periods (**Fig. 6F**, bottom). During the decision and anticipation periods, the vector angle relative to reward value significantly increased (AUC > 0.90), while sum of scalar was not changed (**Fig. 6F**, top). During the outcome period, on the other hand, vector scalar significantly decreased (AUC > 0.90) and vector angle did not change after amygdala lesions. These results suggest that amygdala lesions shifted the population-level neural representation toward interoceptive coding in dACC, whereas coding of all task related information (i.e., both of decision-making and interoceptive coding) generally diminished in OFC.

Next, we conducted the same analysis but, instead of projecting the beta coefficients for HR coding against reward-value coding for each neuron, we projected them against the beta values for choice direction (**Fig. 6G-J**). Here we reasoned that if the impact of increased baseline HR on RTs (**Fig. 2**) is related to changes in dACC then we should expect there to be a change in the vector angle between HR and choice direction encoding. Further, this change should be specific to the decision period of the trial. This is indeed what we observed (**Fig. 6J**, bottom). In dACC there was a change in the vector angle but only in the decision, not anticipation or outcome periods of the trial, indicating an increase in the population-level coding of bodily arousal around the time monkeys were deciding which option to select. In OFC there was only a change in the scalar, similar to what we observed between HR and reward-value encoding (**Fig. 6H**, top).

Despite the population-level change in coding, the correlation between reward-value and HR coding was unchanged by amygdala lesions in both areas (***SI Appendix***, **Fig. S4A**). The correlation between choice direction and HR coding was significantly reduced in OFC during the anticipation period (AUC > 0.90) but not changed in dACC (***SI Appendix*, Fig. S4B**). Thus, changes in population coding were not simply accounted for by the change in correlation between correlation coefficients for either reward value or choice direction and baseline HR. In summary, bilateral amygdala lesions altered the balance of coding between either reward value or choice direction and baseline HR representations in dACC, skewing representations towards the current state of bodily arousal. Thus, our data indicate that hyperarousal alters the relationship between HR and RTs by heightening the impact of bodily arousal on decision-making mechanisms in dACC.

## DISCUSSION

How bodily arousal influences decision-making has been a central question in psychology and neuroscience. What has not been clear is how these processes interact at the level of single neurons. Here we found that higher HR and reward size were associated with faster RTs in macaques performing a reward-guided decision-making task (**Fig. 2**). Concurrently, a population of neurons in OFC and dACC encoded the current HR on each trial (**Figs 3** and **4**). Notably, representations of HR were largely independent of reward value, suggesting that arousal and reward-value signals are represented separately in frontal cortex.

Bilateral excitotoxic lesions of the amygdala produced a tonic increase in bodily arousal and this was associated with a marked change in how HR and reward value impacted RTs (**Fig. 2**). Whereas before amygdala lesions subjects’ behavior was characterized by a negative correlation between arousal and RTs, after amygdala lesions this changed to a positive correlation. Concurrent to the change in bodily arousal and its impact on RTs, the proportion of neurons encoding the current HR increased in dACC, but not OFC (**Fig. 5**). At the population level, increased bodily arousal was associated with a shift in the balance of encoding away from signaling decision-related variables, namely reward value or choice direction, and towards signaling HR in dACC but not OFC (**Fig. 6**). Thus, changes in dACC encoding of the current state of bodily arousal are associated with altered decision-making.

### The influence of bodily arousal on value-based decision-making

Previous work showed that amygdala dysfunction upregulates arousal states in animals and humans (22, 29), but how amygdala-linked changes in bodily arousal alter decision-making have been largely unexplored. We found that, in intact monkeys, higher HR during the first part of each trial, which we take as a measure of current bodily arousal, was negatively correlated with RTs to choose different rewarded options. This indicates that bodily arousal broadly facilitates responding under normal circumstances. Notably, this effect did not interact with the size of reward available on each trial indicating that reward size and bodily arousal have distinct influences on behavior (**Fig. 2**). Although we recorded HR, we likely would have obtained similar results if we had recorded other measures of bodily arousal such as blood pressure, pupil diameter, or skin conductance responses. This is because these autonomic measures are highly correlated (22, 30).

When bodily arousal increased following bilateral excitotoxic lesions of amygdala, we found that the influence of HR on RTs was reversed. We interpret this change in relation to the classic Yerkes-Dodson law (25) whereby increasing levels of arousal become detrimental to performance (24). Here bodily arousal slowed decisions, delaying access to reward. We interpret this effect on behavior in terms of bodily arousal, as opposed to direct task-based influences of amygdala lesions, as HR was elevated even at rest, not just during the task (***SI Appendix***, **Fig. S2**). Thus, the relationship between bodily arousal and behavioral performance followed an inverted ‘U’ function across experiments; amygdala lesions increased the general level of bodily arousal and reversed the impact of bodily arousal on behavioral performance (**Fig. 2**).

Notably, amygdala lesions enhanced the relationship between reward size and RTs. Now, greater amounts of reward had an even stronger negative influence on RTs (**Fig. 2**). Correct performance (accuracy) was not affected by amygdala lesions (***SI Appendix***, **Fig. S1**), but this is likely a ceiling effect as amygdala lesions do impact other types of reward-guided choice behavior (31). The change in the influence of reward value could be related to heightened arousal biasing actions to be more habitual (32, 33), but this latter effect is hard to disentangle from the known influence of amygdala lesions on goal-directed behavior (34). In sum, our findings suggest that, in intact macaques, the amygdala moderates bodily arousal to keep it at a level that promotes efficient performance.

### Frontal cortex representations of bodily arousal and reward

Across an array of decision-making tasks and contexts, neurons within OFC and dACC of macaques signal the magnitude of available rewards and the costs associated with obtaining them (26, 28, 35, 36). Few studies have looked at how bodily arousal is encoded in frontal cortex or might interact with the coding of potential rewards (37). Here we confirm and extend these previous findings by showing that a population of neurons in OFC and dACC encoded the current HR on each trial (**Figs 3** and **4**). Single neuron encoding of bodily arousal in frontal cortex is consistent with findings from neuroimaging studies of humans reporting activations in OFC and ACC (10–13). The neuronal population signaling bodily arousal was found widely across both OFC and dACC, and throughout the whole course of the trial.

We also found that neural correlates of bodily arousal on each trial are largely independent of the available rewards. Note that this finding at the level of single neurons mirrors the separate effects of reward and bodily arousal on behavioral correlates of decision-making (**Fig. 2**). That reward and bodily arousal are represented separately is potentially surprising as larger reward magnitudes invigorate responding, increasing arousal. One possible explanation for this difference is that the amount of reward available on each trial has a transient influence on bodily arousal. This is because the rewards available on each trial are randomly chosen and cue-reward associations were extensively learned before testing. By contrast, HR during fixation period incorporates multiple long-term influences on arousal such as recent history of reward, overall satiety, and fatigue. Note, however, that we did not find an effect of prior reward history on baseline HR suggesting that simple reward-based explanations cannot account for our findings (***SI Appendix***, **Fig. S2**).

### Representations of bodily arousal in dACC and its influence on decision-making

The proportion of neurons encoding HR increased in dACC, but not in OFC, after amygdala lesions, whereas the proportion of reward-value coding neurons preferentially decreased in OFC (**Fig. 5**). Statistically there was no change in the proportion of neurons coding choice direction. The change in HR coding in dACC was evident at the level of the whole population of recorded neurons as a change of vector angle towards signaling HR, while any changes in OFC were solely a reduction of the vector scalar (**Fig. 6**). Note that the enhancement of HR signaling in dACC mainly occurred at the population level rather than at the single-cell level; we did not detect a shift in the proportion of neurons coding the interaction between reward value or choice direction and HR (**Fig. 5**). This result is potentially important as it speaks to predictive coding accounts of interoceptive awareness (10, 38). Here the population-level change of interoceptive representations in dACC could reflect a general change in subjective interoceptive sensitivity, which would then lead to excess mismatch between subjective and objective representations in bodily states, which disrupts decision-making.

The change in coding in dACC suggests that the lesions and their associated increase in HR caused a shift in the balance of encoding in dACC across the whole population, indicative of a change in information processing. That amygdala lesions caused a specific change in the balance of coding between bodily arousal and choice direction during the decision period potentially pinpoints where in time bodily arousal influences RTs in particular and decision-making in general (**Fig. 2**). On this view, increased HR coding in the population of dACC neurons interferes with choice direction coding when animals are choosing between options (**Fig. 6**), slowing RTs in the period most critical for decision-making.

### dACC, visceromotor function, and interoception

There are at least three possible accounts, not mutually exclusive, for the changes in dACC encoding of bodily arousal. The altered encoding could be: 1) simply the direct result of loss of input from the amygdala; 2) related to dACC directly driving higher states of bodily arousal; or 3) indicative of altered interoceptive representations of bodily arousal and/or their integration into decision-making. Fully distinguishing among these alternatives is beyond the scope of the present study. However, we favor the latter two interpretations for the following reasons. First, the part of dACC that we recorded from in the dorsal bank of the cingulate sulcus receives only sparse innervation from amygdala (39). This argues against direct amygdala lesion effects as does the non-task specific impact of amygdala lesions on HR (***SI Appendix***, **Fig. S2B**). Second, dACC has been given a role in both active sensing of interoceptive states as well as controlling autonomic arousal (11), likely through connections to hypothalamus (40, 41). Third, direct stimulation of the dACC in humans and macaques causes changes in heart rate (42, 43). This would appear to argue in favor of increased HR encoding in dACC being the driver of changes in arousal following amygdala lesions, in keeping with dACC as a visceromotor effector brain region. Against this, lesions of dACC in macaques and humans are not always associated with changes in peripheral physiology (11, 44), which argues in favor of dACC representing the current state of bodily arousal. The interoceptive representation in dACC could reflect mechanoreceptive signaling of heart rate that is carried through sympathetic and parasympathetic afferents, although it could reflect other factors, such as thirst being integrated into subjective valuations (10). In the current study, however, we did not find an interaction between baseline HR and reward history (***SI Appendix***, **Fig. S2**), providing more evidence in favor of the idea that the interoceptive representation in dACC are tracking spontaneous fluctuations in HR. Thus, changes in HR coding in dACC may reflect both increased interoceptive representations of bodily arousal as well as visceromotor effectors of bodily arousal.

### Summary

Our results provide unique insights into how bodily arousal impacts reward-guided decision-making at both behavioral and neural levels. Our data point to interoceptive coding in frontal cortex as the mechanism through which bodily arousal impacts decision-making. This mechanism may underlie a wide array of psychological phenomena and potentially explain the cognitive biases observed in certain psychiatric diseases characterized by persistent hyper-arousal (6–8). Although the task used in the current study did not directly test the relationship between choice accuracy and bodily arousal, future studies could tackle this question by using learning or gambling tasks, in which choice options are subjectively valued and therefore choices are likely more susceptible to the interoceptive influence of bodily states.

## METHODS

### Subjects, Task, Surgical procedures, and Neuronal recordings

The full details of the experimental procedures were reported in a previous paper (20). Here, we provide a brief description of the procedures. Three male rhesus macaques (*Macaca mulatta*: monkeys H, N, and V) weighing 8.5, 8.0, and 8.4 kg, respectively, were used in the study. All procedures were reviewed and approved by the National Institute of Mental Health (NIMH) Animal Care and Use Committee. In the two-choice visually guided task, monkeys had to press and hold a central button and then fixate a central red spot for 0.5–1.5 s to begin a trial (**Fig. 1A**). Two visual stimuli (S1 and S2) associated with different amounts of fluid reward (0, 0.1, 0.2, 0.4, or 0.8 ml) were then sequentially presented separated by an interval of 1 second (**Fig. 1B**). The order and position of the stimuli (right or left of the central spot) were randomized. After a variable delay (0.0–1.5 s), the central spot brightened (‘Go’ signal), and the monkeys were required to make a choice by moving their hand to the left or right response button. The amount of reward corresponding to the chosen stimulus was delivered 0.5 s later.

Monkeys were implanted with a head restraint device and a recording chamber over the frontal lobe. After the preoperative recordings were completed, MRI-guided bilateral excitotoxic lesions of the amygdala were made in each monkey. Single-unit recordings were performed with tungsten microelectrodes. All OFC recordings were between 27 and 38 mm anterior to the interaural plane (Walker’s areas 11 and 13). dACC neurons were recorded between the anterior tip of the cingulate sulcus (approximately 38 mm) and 24 mm (areas 9 and 24). The recording sites were verified by MRI with electrodes in place and histological analysis of marking lesions, and the extents of the amygdala lesions were estimated by microscopic examination of Nissl-stained brain sections (20).

### ECG data acquisition and pre-processing

The detailed methods of ECG recording and analysis were described in a previous paper (22). In brief, surface ECG electrodes were placed on the monkey’s back, and leads were placed over the rib-cage and scapula. The HR recording was performed throughout behavioral test sessions, concurrently with single-neuron recording. ECG signals were low-pass filtered at 360 Hz, then digitized at 1000 samples/s. The heart rate was calculated from the interval between R waves identified using custom software specifically designed for macaque ECG (22).

### Data analysis

On each day, single neurons were isolated and activity recorded for >100 trials before electrodes again were moved and another set of neurons isolated. This meant that in a single day of recording there were many separate sessions where neurons were isolated. Only recording sessions in which neuronal data and ECG data were collected simultaneously were analyzed here (pre-op: n = 57, 68, 83 sessions, post-op: n = 31, 130, 52 sessions for monkeys H, N, V, respectively). Within these sessions, analyses focused on ‘correct trials’ where subjects chose the stimulus associated with the greater amount of reward. The reaction time (RT) was defined as the interval from the onset of the ‘Go’ signal until release of the central button. For the heart rate and neuronal analysis, four analysis epochs were defined as follows: “fixation period” (−1000 to 0 ms before S1 onset) in which animals keep fixation and neither option has been presented; “decision period” (0 to 1000 ms after S2 onset) in which both options are revealed and subjects are able to decide which option to choose; “anticipation period” (1000 to 0 ms before reward) where subjects have made a choice and are waiting for reward; and “outcome period” (0 to 1000 ms after reward) in which animals receive the chosen reward amount. Note that the anticipation period will overlap a little with the decision period. Also note that this is different to the 500 ms “expected reward” defined in the previous study (20). Here all periods were set to 1000 ms for consistency.

To examine the change of arousal state outside the task setting, ECG was recorded in ‘resting’ sessions, in which monkeys were not engaged in any task and were sitting quietly in the primate chair while electrodes settled prior to recordings (pre-op: n = 1, 34 sessions for monkeys H and V, post-op: n = 1, 44, 33 sessions for monkeys H, N, V, respectively). The heart rate during resting sessions (resting HR) was computed in the same manner as during the decision-making task, and the impact of bilateral amygdala lesions was assessed by one-way repeated-measures ANOVAs with monkey modeled as a random effect.

The baseline heart rate for each trial (baseline HR) was defined as the heart rate during the fixation period. The continuous value of baseline HR was used for the following analyses. To test the effect of reward size and the amygdala lesions on the baseline HR, we performed two-way repeated-measures ANOVAs. To examine the relationship between the RT, reward size, and heart rate, we performed a multiple linear regression analysis as below:

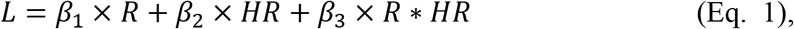

where, *L*, *R*, *HR*, and *R*HR* indicate RT, reward size, baseline HR, and interaction of reward size and baseline HR, respectively.

For the neuronal analyses, only correct trials were analyzed. We classified neurons as task-related based on a multiple-regression analysis, as below;

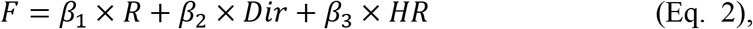

where *F*, *R*, *Dir*, and *HR* mean the firing rate, reward size, choice direction, and the baseline HR, respectively. The neurons that showed significant coding of either the reward size, choice direction, or baseline HR during one of the analysis periods (p < 0.05) were classified as reward-value coding, direction coding, and HR coding neurons, respectively. Significant neurons were further classified as either positive coding (e.g., positive HR coding) or negative coding (e.g., negative HR coding) in accordance with sign of correlation coefficients (*r*) during each analysis periods. We also performed a sliding-window analysis (100-ms bin, 10-ms step) to the task-related neurons using the same linear-regression model.

The population coding analysis was performed using the activity from all recorded neurons. First, we calculated the correlation coefficients for the reward size and baseline HR using the Eq. 2 for each analysis period of every recorded neuron. Separate scatter plots were then made for each area and each analysis period before and after amygdala lesions. The ellipse representing 95% confidence interval and its eigenvectors were drawn on each scatter plot. The population coding was then quantified as the scalar and angle of the eigenvectors; the sum of scalars was computed by adding the scalar of long and short eigenvectors, and the angle to value was computed as the angle from the value-coding to the long eigenvector (**Fig. 6A** and **B**). For the statistical comparison, we computed the vector scalar and angle for randomly sampled populations (80% of data) repeatedly (10,000 times). We then performed ROC analysis and computed the area under the curve (AUC). Our threshold for statistical significance was when the AUC was greater than 0.90.

## Acknowledgments

We are indebted to Vincent Costa for comments on an earlier version of the manuscript. We thank Andrew Mitz, Ravi Chacko, Joshua Ripple, and Kevin Blomstrom for help with data collection. This work was supported by NIMH and the BRAIN initiative (PHR, R01MH110822 and R01MH117040) and by the Intramural Research Program of the National Institute of Mental Health (EAM, ZIAMH002886). AF is also supported by Overseas Research Fellowship from Takeda Science Foundation.

## Author contributions

AF, EAM, and PHR developed the experimental design and ideas for the research; PHR and EAM performed the research; AF and PHR analyzed the data; AF and PHR wrote the manuscript; EAM edited the manuscript.

## Supplementary Information

**Figure S1.**
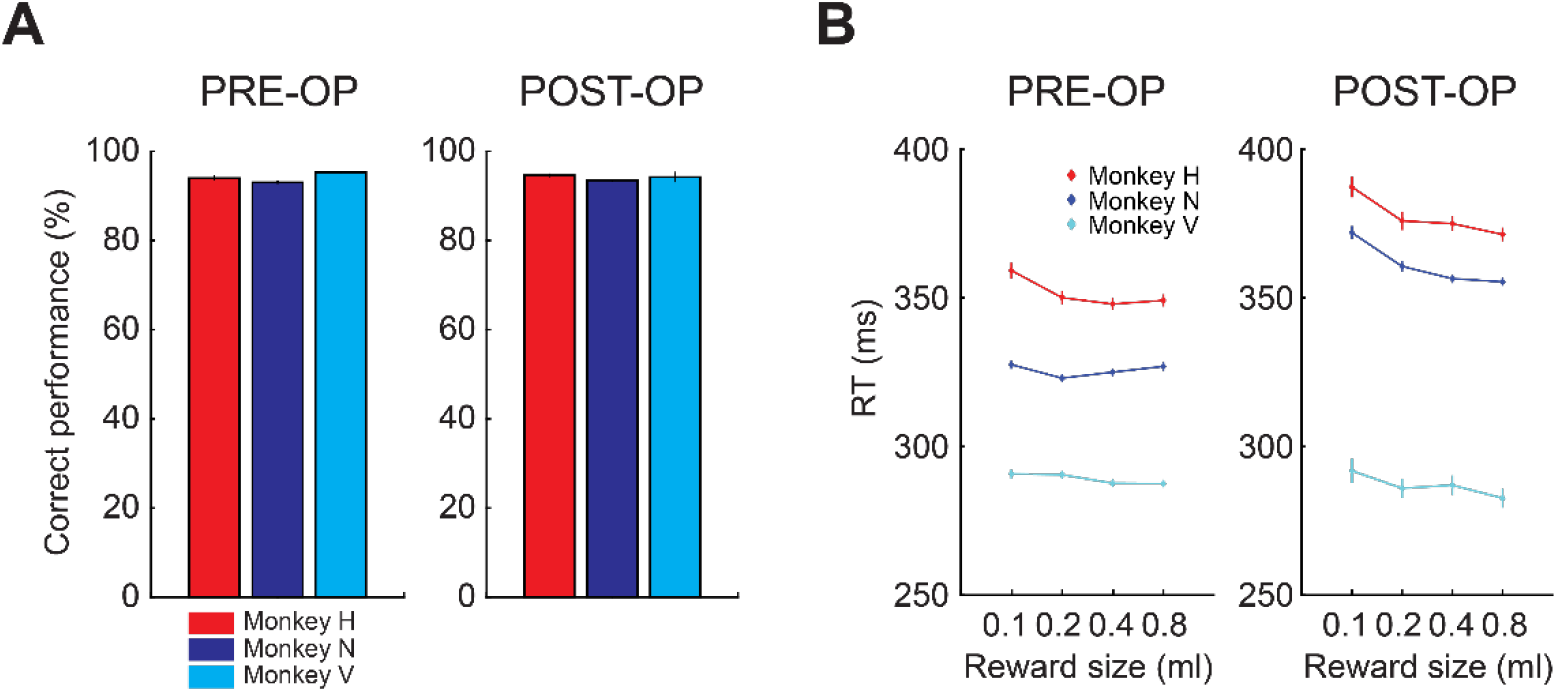
Choice performance in the 2-option reward-guided task. (*A*) Proportion of choices to the stimulus associated with the highest probability of reward on each trial before (left) and after (right) amygdala lesions. (*B*) RTs to choose stimuli associated with 0.1-0.8ml of water reward before (left) and after (right) amygdala lesions. Error bars are standard error of the mean.

**Figure S2.**
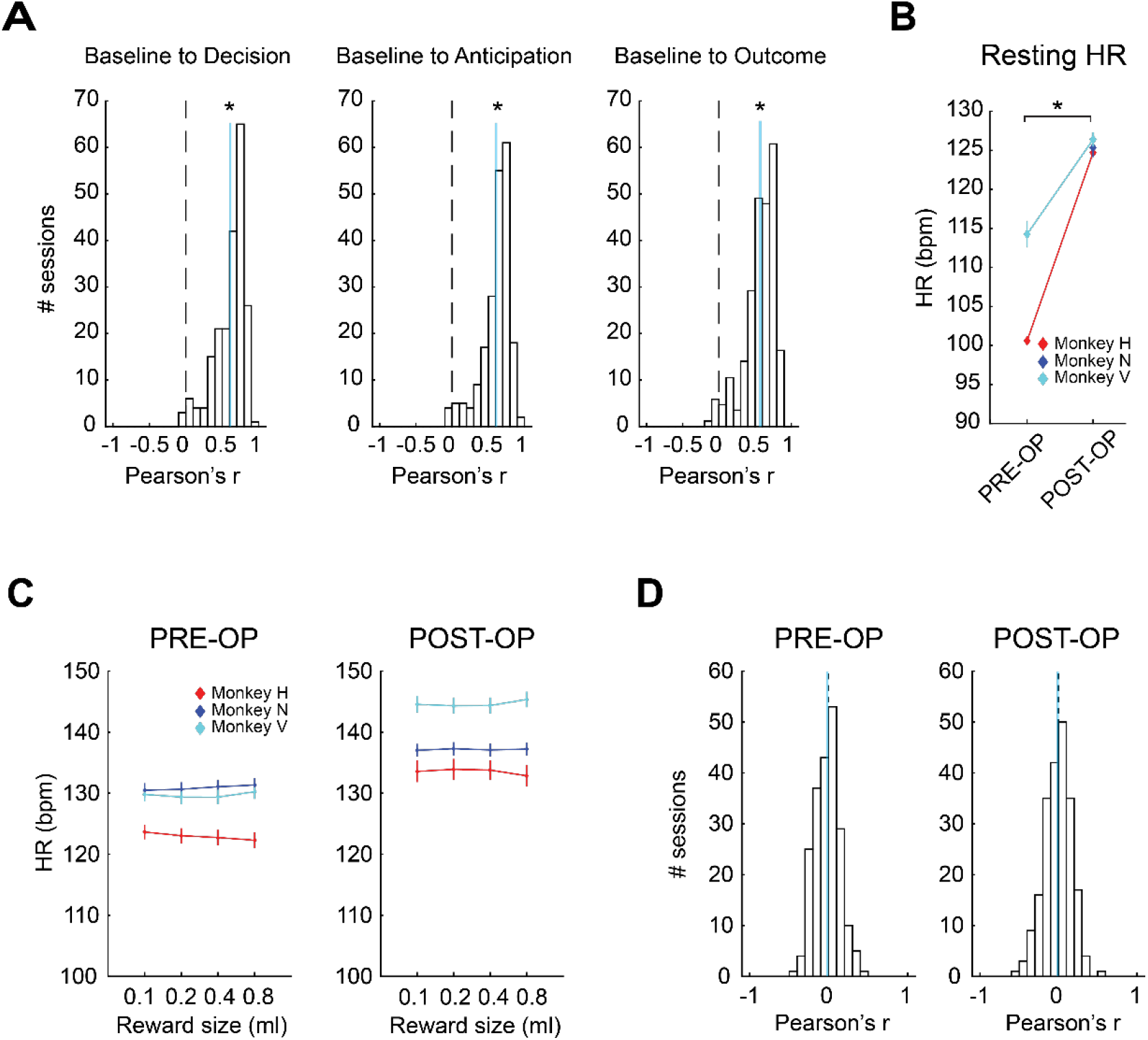
No correlation between reward and baseline HR. (*A*) Distribution of coefficients (*r*) from correlations between the baseline HR and the HR during the decision (left), anticipation (center) and outcome (right) periods, respectively. Cyan lines indicate average *r*. Asterisks indicate p < 0.017 by rank-sum test. (*B*) The heart rate during rest (resting HR) before and after amygdala lesions. During rest, monkeys sat quietly in a primate chair and were not engaged in a behavioral task. For Monkey N (Blue), data was only collected for post-operative sessions. Asterisk indicates p < 0.05 by one-way ANOVA. (*C*) Baseline HR associated with delivery of 0.10.8ml of water reward before (left) and after (right) amygdala lesions. Error bars are standard error of the mean. (*D*) Distribution of correlation coefficients (*r*) that tested correlation between baseline HR and average of earned reward in the preceding 10 trials, before (left) and after (right) amygdala lesions. Cyan lines show average *r*.

**Figure S3.**
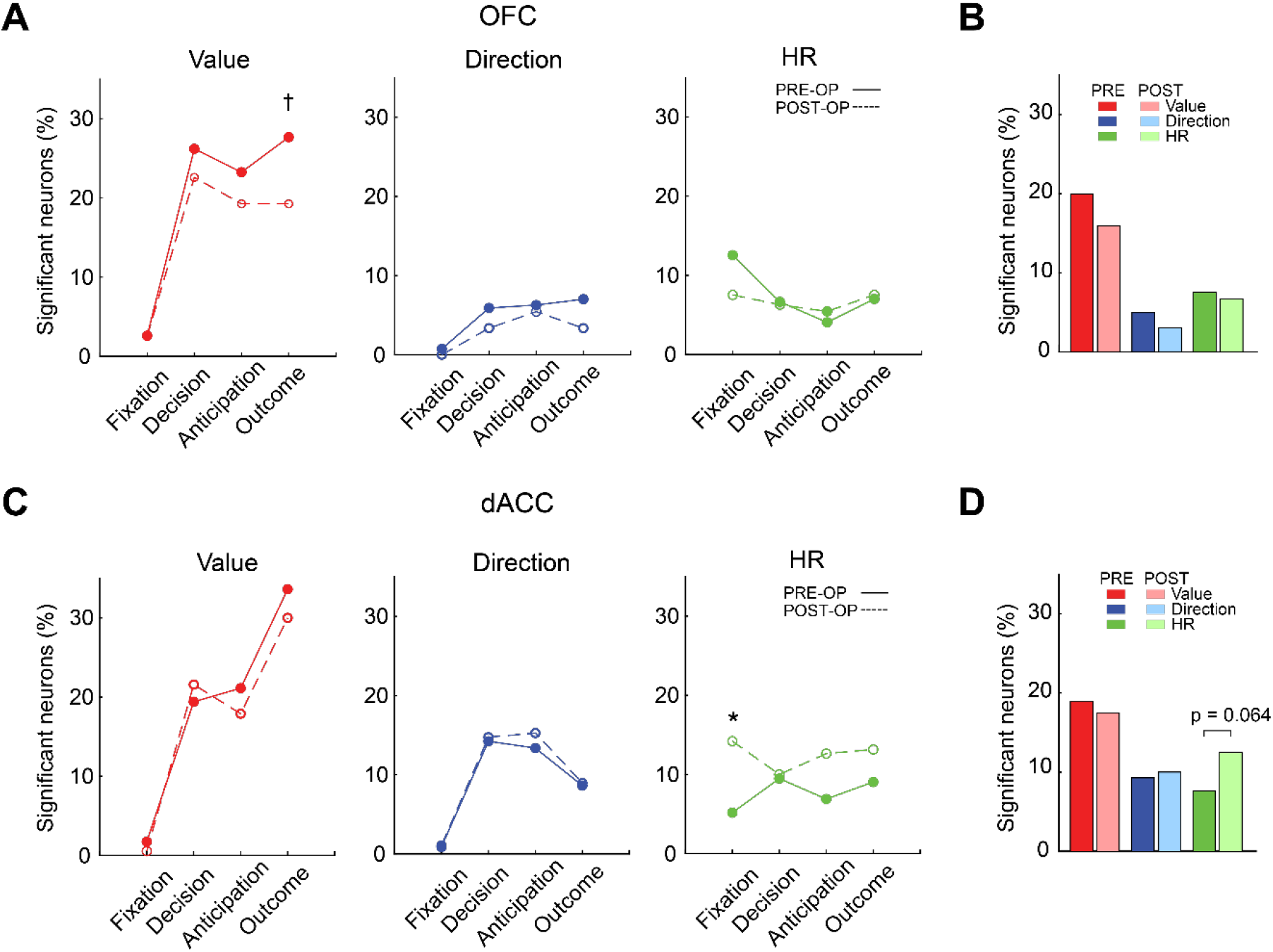
Effect of bilateral amygdala lesions on the proportion of HR coding neurons with corrected threshold. (*A and B*) Proportion of significant reward-value coding, direction coding, and HR coding neuron in OFC. (*C and D*) Proportion of neurons encoding same variables in dACC. Asterisks indicate significant difference in the proportion of neurons (Pre-vs. Post) for each coding type in each analysis period (* p < 0.05, † p < 0.10 with Bonferroni correction, Chi-square test).

**Figure S4.**
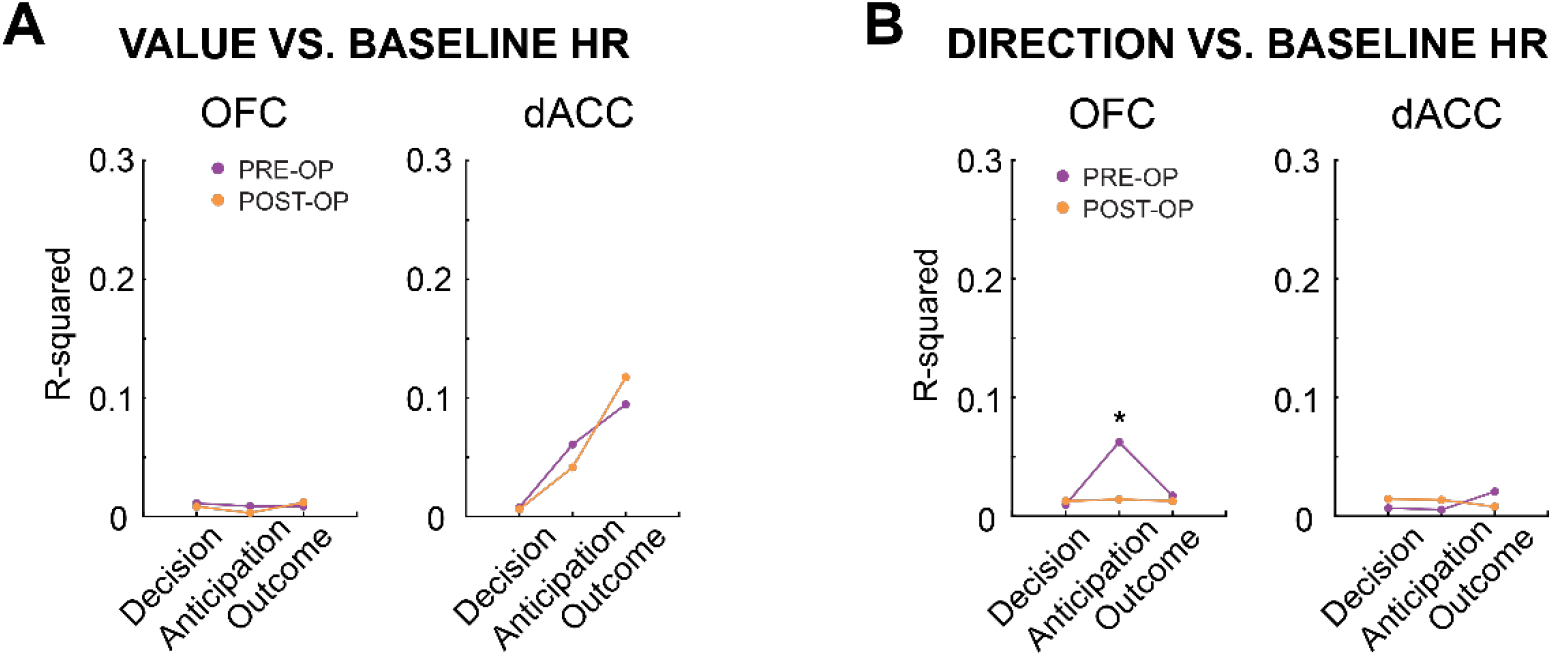
Population coding, correlation analysis. (*A*) Correlation strength (R^2^) between beta for reward-value and beta for baseline HR for each analysis epoch are plotted for OFC (left) and dACC (right), respectively. (*B*) R^2^ value between beta for choice direction and baseline HR. Asterisk indicates AUC > 0.90.

**Table S1.** The number of neurons analyzed and well-isolated single neurons (within brackets) for each monkey.

| *Monkey* | *Treatment* |  |  |  |
| --- | --- | --- | --- | --- |
|  | *Pre-op* |  | *Post-op* |  |
|  | *Area* |  | *Area* |  |
|  | *OFC* | *dACC* | *OFC* | *dACC* |
| *H* | 31 (28) | 102 (97) | - | 83 (77) |
| *N* | 105 (84) | 117 (98) | 142 (128) | 107 (89) |
| *V* | 135 (133) | 13 (11) | 97 (95) | - |
| *Total* | 271 (245) | 232 (206) | 239 (233) | 190 (166) |

**Table S2.** The number of significant neurons in each analysis epoch.

| *Analysis epoch* | *OFC (n = 271)* |  |  | *dACC (n = 232)* |  |  |
| --- | --- | --- | --- | --- | --- | --- |
|  | *Neuron classification* |  |  | *Neuron classification* |  |  |
|  | *Val* | *Direc* | *HR* | *Val* | *Direc* | *HR* |
| *Fixation* | 20 | 8 | 57 | 16 | 11 | 25 |
| *Decision* | 96 | 29 | 42 | 63 | 51 | 36 |
| *Anticipation* | 84 | 29 | 26 | 67 | 46 | 34 |
| *Outcome* | 102 | 34 | 44 | 103 | 36 | 39 |

Val: reward-value coding, Direc: choice direction coding, HR: HR coding.

## REFERENCES

1. Lambourne K & Tomporowski P (2010) The effect of exercise-induced arousal on cognitive task performance: a meta-regression analysis. Brain research 1341:12–24.

2. Wilson M & Daly M (2004) Do pretty women inspire men to discount the future? Proceedings. Biological sciences 271 Suppl 4(Suppl 4):S177–179.

3. Mobbs D & Kim JJ (2015) Neuroethological studies of fear, anxiety, and risky decision-making in rodents and humans. Current opinion in behavioral sciences 5:8–15.

4. Fanselow MS & Lester LS (1988) A functional behavioristic approach to aversively motivated behavior: Predatory imminence as a determinant of the topography of defensive behavior.

5. Mobbs D, et al. (2009) From threat to fear: the neural organization of defensive fear systems in humans. The Journal of neuroscience: the official journal of the Society for Neuroscience 29(39):12236–12243.

6. Herman AM, Critchley HD, & Duka T (2018) The role of emotions and physiological arousal in modulating impulsive behaviour. Biological psychology 133:30–43.

7. Paulus MP & Stewart JL (2014) Interoception and drug addiction. Neuropharmacology 76 Pt B(0 0):342–350.

8. White SF, et al. (2017) Prediction Error Representation in Individuals With Generalized Anxiety Disorder During Passive Avoidance. The American journal of psychiatry 174(2):110–117.

9. Craig AD (2002) How do you feel? Interoception: the sense of the physiological condition of the body. Nat Rev Neurosci 3(8):655–666.

10. Craig AD (2009) How do you feel--now? The anterior insula and human awareness. Nature reviews. Neuroscience 10(1):59–70.

11. Critchley HD, et al. (2003) Human cingulate cortex and autonomic control: converging neuroimaging and clinical evidence. Brain: a journal of neurology 126(Pt 10):2139–2152.

12. Critchley HD, Wiens S, Rotshtein P, Ohman A, & Dolan RJ (2004) Neural systems supporting interoceptive awareness. Nature neuroscience 7(2):189–195.

13. Wang J, et al. (2005) Perfusion functional MRI reveals cerebral blood flow pattern under psychological stress. Proceedings of the National Academy of Sciences of the United States of America 102(49):17804–17809.

14. Rushworth MF, Noonan MP, Boorman ED, Walton ME, & Behrens TE (2011) Frontal cortex and reward-guided learning and decision-making. Neuron 70(6):1054–1069.

15. Bechara A, Damasio H, & Damasio AR (2000) Emotion, decision making and the orbitofrontal cortex. Cerebral cortex (New York, N.Y.: 1991) 10(3):295–307.

16. Bechara A, Damasio H, Damasio AR, & Lee GP (1999) Different contributions of the human amygdala and ventromedial prefrontal cortex to decision-making. The Journal of neuroscience: the official journal of the Society for Neuroscience 19(13):5473–5481.

17. Bechara A, Damasio H, Tranel D, & Damasio AR (1997) Deciding advantageously before knowing the advantageous strategy. Science (New York, N.Y.) 275(5304):1293–1295.

18. Rudebeck PH, et al. (2014) A role for primate subgenual cingulate cortex in sustaining autonomic arousal. Proceedings of the National Academy of Sciences of the United States of America 111(14):5391–5396.

19. Mobbs D, Headley DB, Ding W, & Dayan P (2020) Space, Time, and Fear: Survival Computations along Defensive Circuits. Trends in cognitive sciences 24(3):228–241.

20. Rudebeck PH, Mitz AR, Chacko RV, & Murray EA (2013) Effects of amygdala lesions on reward-value coding in orbital and medial prefrontal cortex. Neuron 80(6):1519–1531.

21. McNamara L & Ballard ME (1999) Resting arousal, sensation seeking, and music preference. Genetic, Social, and General Psychology Monographs 125(3):229.

22. Mitz AR, Chacko RV, Putnam PT, Rudebeck PH, & Murray EA (2017) Using pupil size and heart rate to infer affective states during behavioral neurophysiology and neuropsychology experiments. J Neurosci Methods 279:1–12.

23. Clynes M (1960) Computer analysis of reflex control and organization: respiratory sinus arrhythmia. Science (New York, N.Y.) 131(3396):300–302.

24. Hebb DO (1955) Drives and the C.N.S. (conceptual nervous system). Psychological review 62(4):243–254.

25. Yerkes RM & Dodson JD (1908) The relation of strength of stimulus to rapidity of habit - formation. Journal of comparative neurology and psychology 18(5):459–482.

26. Kennerley SW & Wallis JD (2009) Encoding of reward and space during a working memory task in the orbitofrontal cortex and anterior cingulate sulcus. Journal of neurophysiology 102(6):3352–3364.

27. Padoa-Schioppa C & Assad JA (2006) Neurons in the orbitofrontal cortex encode economic value. Nature 441(7090):223–226.

28. Thorpe SJ, Rolls ET, & Maddison S (1983) The orbitofrontal cortex: neuronal activity in the behaving monkey. Experimental brain research 49(1):93–115.

29. Quirk GJ & Gehlert DR (2003) Inhibition of the amygdala: key to pathological states? Annals of the New York Academy of Sciences 985:263–272.

30. Bradley MM, Miccoli L, Escrig MA, & Lang PJ (2008) The pupil as a measure of emotional arousal and autonomic activation. Psychophysiology 45(4):602–607.

31. Rudebeck PH, Ripple JA, Mitz AR, Averbeck BB, & Murray EA (2017) Amygdala Contributions to Stimulus-Reward Encoding in the Macaque Medial and Orbital Frontal Cortex during Learning. J Neurosci 37(8):2186–2202.

32. Schwabe L & Wolf OT (2013) Stress and multiple memory systems: from ‘thinking’ to ‘doing’. Trends in cognitive sciences 17(2):60–68.

33. Balleine BW & Dickinson A (1998) Goal-directed instrumental action: contingency and incentive learning and their cortical substrates. Neuropharmacology 37(4-5):407–419.

34. Málková L, Gaffan D, & Murray EA (1997) Excitotoxic lesions of the amygdala fail to produce impairment in visual learning for auditory secondary reinforcement but interfere with reinforcer devaluation effects in rhesus monkeys. The Journal of neuroscience: the official journal of the Society for Neuroscience 17(15):6011–6020.

35. Cai X & Padoa-Schioppa C (2012) Neuronal encoding of subjective value in dorsal and ventral anterior cingulate cortex. The Journal of neuroscience: the official journal of the Society for Neuroscience 32(11):3791–3808.

36. Shima K & Tanji J (1998) Role for cingulate motor area cells in voluntary movement selection based on reward. Science (New York, NY) 282(5392):1335–1338.

37. Ebitz RB & Platt ML (2015) Neuronal activity in primate dorsal anterior cingulate cortex signals task conflict and predicts adjustments in pupil-linked arousal. Neuron 85(3):628–640.

38. Critchley HD & Garfinkel SN (2017) Interoception and emotion. Current opinion in psychology 17:7–14.

39. Ghashghaei HT, Hilgetag CC, & Barbas H (2007) Sequence of information processing for emotions based on the anatomic dialogue between prefrontal cortex and amygdala. NeuroImage 34(3):905–923.

40. Ongür D, An X, & Price JL (1998) Prefrontal cortical projections to the hypothalamus in macaque monkeys. The Journal of comparative neurology 401(4):480–505.

41. Dum RP, Levinthal DJ, & Strick PL (2019) The mind-body problem: Circuits that link the cerebral cortex to the adrenal medulla. Proceedings of the National Academy of Sciences of the United States of America 116(52):26321–26328.

42. Kaada BR, Pribram KH, & Epstein JA (1949) Respiratory and vascular responses in monkeys from temporal pole, insula, orbital surface and cingulate gyrus; a preliminary report. Journal of neurophysiology 12(5):347–356.

43. Lewin W & Whitty CW (1960) Effects of anterior cingulate stimulation in conscious human subjects. Journal of neurophysiology 23:445–447.

44. Basile BM, et al. (2020) Autonomic arousal tracks outcome salience not valence in monkeys making social decisions. bioRxiv.2020.2002.2017.952986.

